# Transcript isoform differences across human tissues are predominantly driven by alternative start and termination sites of transcription

**DOI:** 10.1101/127894

**Authors:** Alejandro Reyes, Wolfgang Huber

## Abstract

Most human genes have multiple transcription start and polyadenylation sites, as well as alternatively spliced exons. Although such transcript isoform diversity contributes to the differentiation between cell types, the importance of contributions from the different isoform generating processes is unclear. To address this question, we used 798 samples from the Genotype-Tissue Expression (*GTEx*) to investigate cell type dependent differences in exon usage of over 18,000 protein-coding genes in 23 cell types. We found tissue-dependent isoform usage in about half of expressed genes. Overall, tissue-dependent splicing accounted only for a minority of tissue-dependent exon usage, most of which was consistent with alternative transcription start and termination sites. We verified this result on a second, independent dataset, Cap Analysis of Gene Expression (CAGE) data from the FANTOM consortium, which confirmed widespread tissue-dependent usage of alternative transcription start sites. Our analysis identifies transcription start and termination sites as the principal drivers of isoform diversity across tissues. Moreover, our results indicate that most tissue-dependent splicing involves untranslated exons and therefore may not have consequences at the proteome level.

## Introduction

Alternative splicing, alternative promoter usage and alternative polyadenylation are interdependent molecular processes that enable genomes to synthesize multiple transcript isoforms per gene^1^. In mammalian genomes, it has been estimated that at least 70% of genes have multiple polyadenylation sites, more than 50% of genes have alternative transcription start sites and nearly all genes undergo alternative splicing^2,3,4^. Hence, these molecular processes increase the repertoire of transcripts in mammalian genomes^5,6,7^.

Important biological processes are regulated by the expression of alternative isoforms^8,9^, and their mis-regulation has been observed in many diseases, including cancer^10,11,12,13^. For dozens of genes, it has been experimentally demonstrated that alternative isoforms result in proteins with differences in cellular localization, stability, DNA binding properties, lipid binding properties or enzymatic activity^14^. Alternative isoforms can behave like completely distinct proteins when considering their protein-protein interaction capabilities^15^. Recently, it has been reported that a large majority of alternatively spliced RNAs bind to ribosomes^16^, which suggests that most of them could be translated into proteins and that the currently known instances of functional protein isoforms are only the tip of the iceberg.

On the other hand, based on evidence such as the conservation of protein structures and functional features, it has been concluded that the majority of alternative transcript isoforms would translate into proteins with disrupted structures and functions^17,18^. Indeed, large-scale proteomics surveys indicate that the abundance of such ‘non-functional proteins’, if not zero, is generally below levels that can currently be detected with high confidence^19,20^. This raises the possibility that the functions of a large proportion of transcript isoforms is on the level of the RNA rather than the protein.

If alternative transcript isoforms are functional primarily at the mRNA level, one might expect an important role of alternative transcription start and stop sites, since these have been shown to often enhance post-transcriptional regulation by fine-tuning the stability and translation of mRNAs^21,22,23,24^. Larger contributions of alternative transcription start and stop sites to isoform diversity, as compared to alternative splicing, have been reported in an analysis of transcript databases^25^ and in an analysis of mouse cerebellar development^26^. However, the extent to which each of these isoform generating processes result in isoform differences across many human cell types and contribute to cellular phenotypes is currently unknown.

Here, we developed an analytic strategy to approach these questions using a unique, large data resource established by the Genotype-Tissue Expression (*GTex*) Project V6^27^. We show that alternative transcript isoform choice is specifically regulated across tissues for a large part of the human genome, affecting about half of multi-exonic genes. The majority of these events cannot be explained by alternative splicing; rather, most appear due to alternative usage of start and termination sites of transcription. Integration of data from the *FANTOM* consortium^6^ confirms prevalent tissue-dependent usage of alternative transcription start sites. Our results also suggest that for the majority of genes, alternative splicing has more effects on the RNA rather than the protein product and that alternative transcript start and polyadenylation sites play a large, and previously underexplored role in establising cell type specificity.

## RESULTS

### The data

We used transcriptome data (RNA-seq) from the *V6* release of the Genotype-Tissue Expression (*GTEx*) project^27^. The dataset comprised 9,795 RNA-seq samples from 54 tissues out of a total of 551 human individuals. Since the dataset did not contain each tissue for each individual, we identified subsets of data that could be analyzed as fully crossed designs (i.e., contained all possible tissue–individual combinations). We mapped the sequenced fragments to the human reference genome version *GRCh38*, obtained from *ENSEMBL* release 84^28^, using the aligner *STAR v2.4.2a*^29^. We excluded samples where the number of sequenced fragments was below 1,000,000 or where the percentage of uniquely mapping reads was below 60%. Using these filtering criteria, we defined three subsets of *GTEx* data:

- *Subset A* consisted of 8 brain cell types (frontal cortex [BA9], nucleus accumbens, putamen, cortex, cerebellum, caudate, cerebellar hemisphere and hippocampus) across 30 individuals, comprising a total of 240 samples;
- *Subset B* included 9 tissues (skeletal muscle, thyroid, whole blood, lung, subcutaneous adipose, skin, tibial artery, tibial nerve and esophagus [mucosa]) from 34 individuals, i.e., a total of 306 samples;
- *Subset C* comprised 6 tissues (heart, aorta, esophagus [muscularis], colon and stomach) from 42 individuals (252 samples).

Subsets A, B and C were non-overlapping, and altogether our analysis employed a subset of 798 samples from the *GTEx* dataset.

### Exon usage coefficients enable the study of transcript isoform regulation across tissues

Next, we used *ENSEMBL* transcript annotations to define reduced gene models with non-overlapping exonic regions^30^ (Methods). We obtained 499,667 non-overlapping exonic regions from the 35,048 multi-exonic genes, of which 412,116 were derived from 18,295 protein-coding genes. For each of the subsets A, B and C, we computed two measures of exon usage per exonic region: relative exon usage coefficients (*REUCs*)^31^ and relative spliced-in coefficients (*RSICs*)^4^. The computation of these coefficients is illustrated in Figure 1. Both of them are measures of exon usage in a specific tissue in a particular individual relative to the average exon usage across all tissues and individuals (Methods). The *REUC* defines exon usage as the fraction of sequenced fragments that map to the exonic region among all fragments mapping to the rest of the exonic regions from the same gene. In contrast, the *RSIC* measures the fraction of sequenced fragments that map to the exonic region compared to the number of reads that support the skipping of that exonic region via alternative splicing (Figure 1C). Note that differences in exon usage due to alternative splicing are reflected in both *REUCs* and *RSICs* (Figure 1B and Figure 1C). Changes in exon usage due to alternative transcription initiation sites or alternative polyadenylation sites, which do not result in exon-exon junction reads, are only reflected in *REUCs* (Figure 1D).

**Figure 1:**
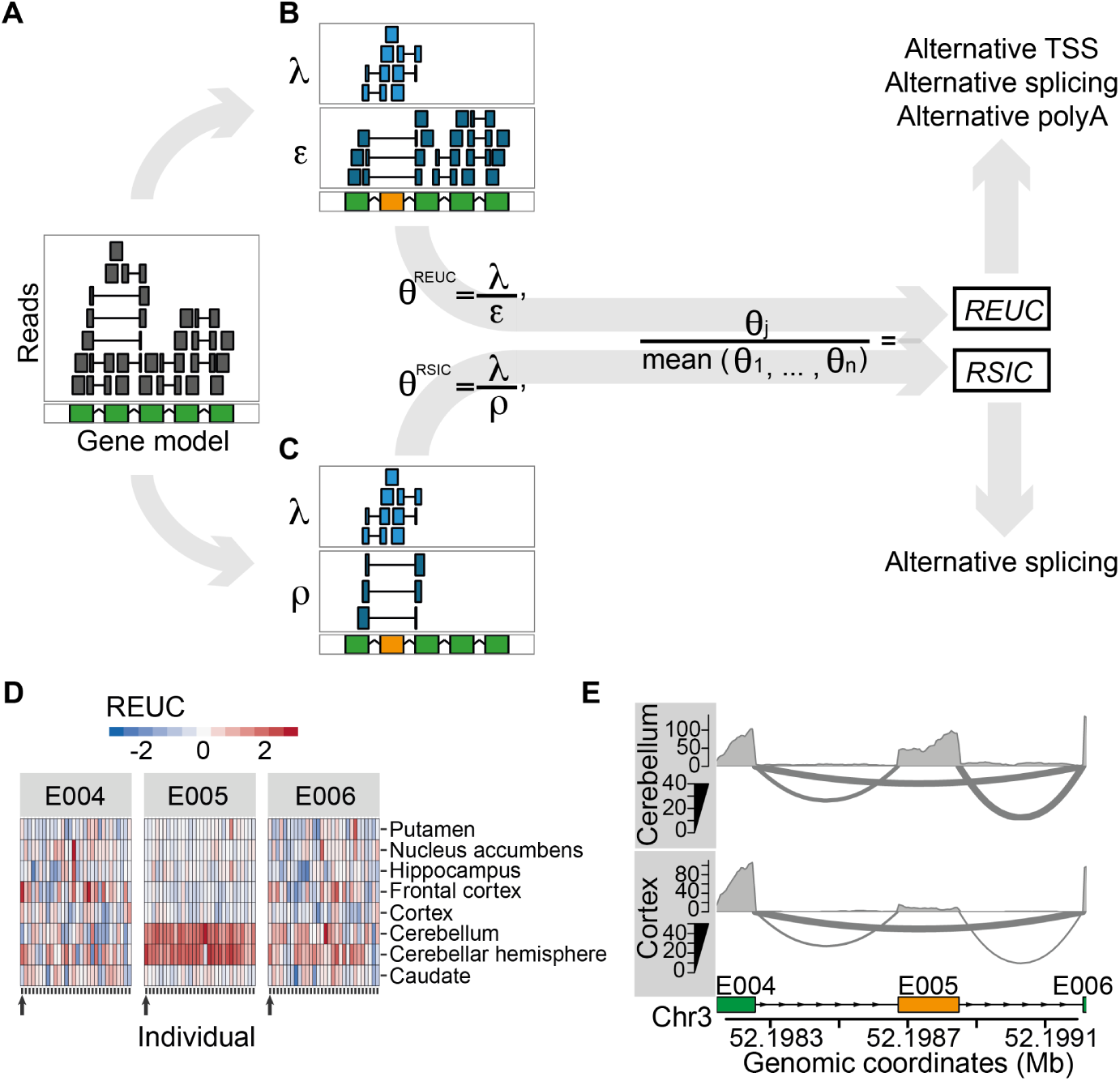
Quantification of exon usage. **(A)** Exemplary gene model in the reference genome (green) and alignments of RNA-seq reads (upper panel). Sequenced fragments whose alignments fall fully into an exonic region are shown by a grey box; alignments that map into two (or possibly more) exonic regions are shown by shorter grey boxes connected by a horizontal line. For a particular exon (highlighted in orange), we consider two strategies to quantify its usage that are illustrated in Panels **B** and **C** (see Methods for the formal description). The first strategy is illustrated in Panel **B**, where sequenced fragments are counted into two groups: the number of sequenced fragments that map fully or partially to the exon (*λ*) and the number of fragments that map to the rest of the exons (*ϵ*). *θ*^REUC^ is defined as the ratio between *λ* and *ϵ*, and the relative exon usage coefficient (*REUC*) for the exon in sample *j* is estimated as the ratio between *θ*^REUC^ in sample *j* with respect to the mean *θ*^REUC^ across the *n* samples. Panel **C** illustrates the second exon usage quantification strategy, where sequenced fragments are also counted into two groups: the number of sequenced fragments that map fully or partially to the exon (*λ*) and the number of sequenced fragments that align to exons both downstream and upstream of the exon under consideration (*ρ*). The latter group of sequenced fragments are derived from transcripts from which the exon was spliced out. *θ*^RSIC^ is now defined as the ratio between *λ* and *ρ*. The relative spliced-in coefficient (*RSIC*) for the exon in sample *j* is estimated as the ratio between *θ*^RSIC^ in sample *j* with respect to the mean *θ*^RSIC^ across the *n* samples. Notice that while differences in exon usage due to alternative splicing are reflected in both *REUCs* and *RSICs*, differences in exon usage due to alternative transcription or termination sites are only reflected in *REUCs*. **(D)** Heatmap representations of the *REUCs* for three exonic regions (*E004*, *E005* and *E006*) of the gene *ALAS1*, computed using *Subset A* of the *GTEx* data. The rows of the heatmaps correspond to the eight tissues, and each column corresponds to one individual. The horizontal colour patterns of exon *E005* indicate elevated inclusion of cerebellum and cerebellar cortex as compared to the rest of the brain cell types. **(E)** RNA-seq samples from two cell types (cortex and cerebellum) from individual *12ZZX* (also indicated by the arrows below each heatmap in Figure 1D) are displayed as sashimi plots^35^. The three exonic regions presented in Panel D are shown. The middle exon, *E005*, is an untranslated cassette exon (ENSEMBL identifier ENSE00002267562) that is spliced out more frequently in cortex than in cerebellum.

As an example, we show the gene *5-Aminolevulinate Synthase 1* (*ALAS1*) in Figure 1D-E. *ALAS1* encodes an enzyme that performs the first catalytic reaction of the biosynthetic pathway of heme. Heme is an organic cofactor that is essential for the proper function and differentiation of many cell types, including those of the hematopoietic, hepatic and nervous systems^32^. Induction of *ALAS1* has been associated with acute attacks of porphyria disease^33^. By exploring the *REUCs* for *ALAS1*, we found that a 5’ untranslated exon was included more frequently in the transcripts generated from cerebellum and cerebellar cortex than in the other brain cell types (Figure 1D). The same pattern of tissue-dependent usage (*TDU*) was also evident from the *RSICs* (Figure S1), which indicates that the *TDU* pattern is a consequence of alternative splicing rather than of alternative transcriptional initiation or termination sites (Figure 1E). In cultured cells, it has been shown that *ALAS1* transcripts that include this 5’ exon are resistant to heme-mediated decay and that their translation is inhibited^34^. The detected splicing pattern indicates that *ALAS1* might be post-transcriptionally regulated differently in cerebellum than in the rest of the brain.

### Tissue-dependent usage of exons is widespread in humans

After observing further instances of tissue-dependent exon usage as in the *ALAS1* example, we asked how common this phenomenon was across the human genome. We defined a score based on the *REUCs* that measures the strength of tissue-dependent expression of an exonic region (Methods). Based on this, we considered an exonic region to be tissue-dependent if its differential usage pattern was statistically significant at a false discovery rate (FDR) of 10%, according to the *DEXSeq* method^30^ and if it had a score larger than 1. We thus detected tissue-dependent usage (TDU) in at least one of the subsets of the *GTEx* data for 23% of the exonic regions (116,601 out of 524,219; Figure S3). Specifically, in *subset A* we retrieved 47,659 exonic regions of 9,839 genes with TDU, in *subset B* 76,562 exonic regions of 12,295 genes, and in *subset C* 30,719 exonic regions of 7,025 genes (Figure 2A and Figure S2). Remarkably, a large fraction of genes detected as expressed were subject to transcript isoform regulation across tissues (Table S1). For example, out of the 18,805 multi-exonic genes that had an average of sequenced fragments of at least 10 in *subset A*, 51% (9,672) showed differential usage in at least one exonic region. Furthermore, the set of genes with TDU was enriched for protein-coding genes as compared to a background set of genes matched for expression strength and number of exonic regions (*p*-value < 2.2·10^−16^, odds-ratio = 3.4, Table S2).

**Figure 2:**
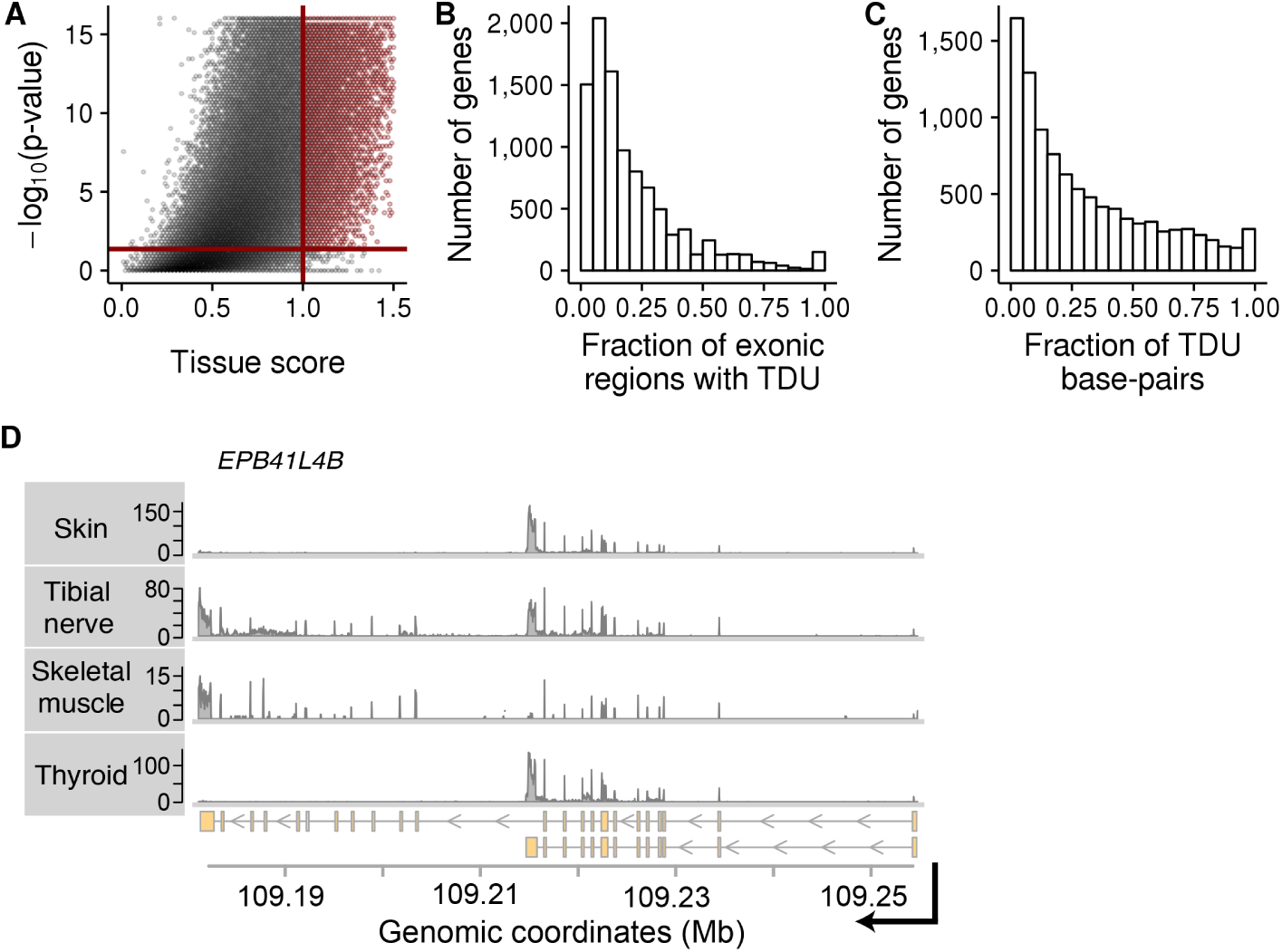
Tissue-dependent exon usage is widespread in the human genome. Panels **A**, **B** and **C** show data from *subset A* of the *GTEx* data. The same plots using data from subsets *B* and *C* can be found in Figure S2 and Figure S4. **(A)** Similar to a volcano-plot, this figure shows statistical significance (p-value on −log_10_ scale) versus effect size (tissue score) of our tissue-dependence test for each exonic region of the human genome. The solid red lines show the thresholds used in this study to call an exonic region tissue-dependent. The p-value threshold 4.28·10^−2^ corresponds to an *adjusted p*-value of 0.1 according to the Benj amini-Hochberg method to control false discovery rate (FDR) for all p-values shown. **(B)** Histogram of the fraction of exonic regions within each gene that are subject to tissue-dependent usage (*x*-axis). The *y*-axis shows the number of genes. **(C)** Similar to Panel B, but expressed in terms of fraction of base-pairs within a gene affected by tissue-dependent usage. **(D)** Exemplary data from four tissues of individual *131XE*. Shown is RNA-Seq coverage (*y*-axis) plots along genomic coordinates (*x*-axis) of chromosome 9 in the locus of the gene *EBP41L4B*. The lower panel shows the transcript annotations for the gene *EPB41L4B*. Skin and thyroid express short isoforms of *EPB41L4B*, while tibial nerve and skeletal muscle express longer isoforms.

We next investigated the nature of transcript isoform differences between tissues. For each gene containing exons with TDU, we estimated the fraction of exonic regions that were subject to TDU and the fraction of affected exonic nucleotides. TDU effects were localized to a relatively small fraction of exons for most genes (Figure 2B, Figure 2C and Figure S4). For instance, in *subset A* the percentage of exonic regions with TDU was below 25% for 70% (6,929) of the genes. In the same subset, the percentage of nucleotides affected by TDU was below 25% for 53% (5,248) of the genes (Table S3). The rest of the cases, where a larger fraction of the gene length was used in a tissue-dependent manner, were instances in which very different isoforms were expressed by different tissues. For example, all27 exonic regions of the gene *Erythrocyte Membrane Protein Band 4.1 Like 4B* (*EPB41 L4B*) were detected to be used in a tissue-dependent manner in *subset B* of the *GTEx* data: in tibial nerve and skeletal muscle, the first 16 exonic regions (counting from 5’ to 3’) had lower usage while the exonic regions located towards the end of the gene showed increased usage (Figure S5 and Figure S6). In this case, the observed pattern can be explained by the two annotated transcript isoforms of the gene: whereas most cell types in *subset B* tend to express the short isoform, tibial nerve and skeletal muscle express the longer isoform more frequently (Figure 2D).

Our results indicate that differences in transcript isoform regulation across tissues is a widespread phenomenon across the human genome. It preferentially affects protein-coding genes. For hundreds of genes, different tissues express very distinct transcript isoforms at different levels.

### Alternative transcriptional initiation and termination sites drive most transcript isoform differences between tissues

In the previous section we saw the example of *EPB41L4B* for a transcript isoform switch that is not driven by alternative splicing, and instead, by the usage of an alternative polyadenylation site (here also referred to as transcription termination site) (Figure 2D). We asked more globally: among the thousands of instances of tissue-dependent exon usage, what fraction is driven by alternative splicing and what by alternative transcriptional start or termination sites?

For each exonic region, we searched for evidence of alternative splicing by counting, in each sample, the number of sequenced fragments that supported exon skipping (as sketched in Figure 1C). We found that only a minor fraction of exonic regions with tissue-dependent usage (TDU) had appreciable evidence of being spliced out from transcripts (Table S4). For instance, the mean of read counts supporting exon skipping was larger than 10 in only 30% (9,282) of the exonic regions with TDU in *subset C*. On the other hand, 58% (17,815) showed no or only weak evidence of being alternatively spliced (Figure 3A and Figure S7). We estimated that alternative splicing explains tissue-dependent transcript differences for at most 36% of the genes (Table S5).

**Figure 3:**
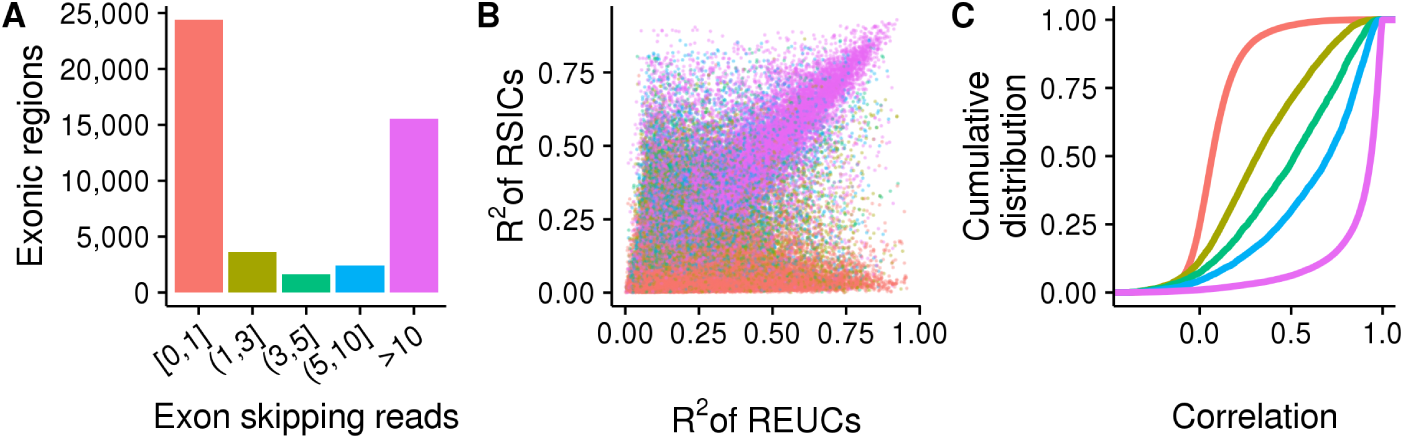
Alternative splicing underlies only a minor fraction of exons with tissue-dependent usage, while the rest are consistent with alternative transcription start or stop sites. The three panels show data from *subset A* of the *GTEx* data. Analogous plots using data from subsets *B* and *C* are shown in Figure S7. **(A)** The heights of the bars show the number of exonic regions with TDU, grouped according to the number of reads that support their splicing out from transcripts. Most exonic regions with TDU have either no or weak evidence of being spliced out from transcripts (bar colored in pink salmon). The bar colors serve also as color legends for Figure 3B and Figure 3C. **(B)** Each point represents one of the 47,659 exonic regions that were detected to be used in a tissue-dependent manner. The x-axis shows the fraction of *REUC* variance that is attributed to variance between tissues (*R*^2^). Analogously, the *y*-axis shows the *R*^2^ statistic for the *RSICs*. Exonic regions with strong evidence of being spliced out from transcripts (purple points) lay along the diagonal. **(C)** Cumulative distribution functions of the Pearson correlation coefficients between the REUCs and the RSICs are shown for exonic regions with TDU. The regions are stratified according to the number of sequenced fragments supporting their splicing out from transcripts. The *REUCs* and *RISCs* are highly correlated for the minor fraction of exons that have strong evidence of being spliced out from transcripts (purple line).

As a second line of evidence, we quantitatively compared the relative exon usage and spliced-in coefficients (*REUCs* and *RSICs*, as defined above). For each exonic region and each subset of the *GTEx* data, we fit two analysis-of-variance models, one for the *REUCs* and one for the *RSICs*, using tissues and individuals as categorical covariates. We determined the coefficient of partial determination (*R*^2^) of the tissue covariate for each fit. A large value of *R*^2^ in the *RSIC* fit indicates that the TDU is due to alternative splicing. Conversly, a large *R*^2^ in the *REUC* fit indicates that the TDU is due to any of alternative splicing, alternative transcription initiation sites or alternative transcriptional termination sites. For the minority of exonic regions with TDU that also had strong evidence of alternative splicing, the *REUCs* and the *RSICs* were highly correlated, and their *R*^2^ statistics were in good agreement, confirming that the TDU was due to alternative splicing (Figure 3B, Figure 3C and Figure S7). Nevertheless, for the majority of exonic regions with TDU, the TDU was consistent with alternative transcription initiation and termination sites.

### Analysis of CAGE data confirms prevalent tissue-dependent usage of alternative transcription start sites

To further investigate the hypothesis that alternative splicing does not drive most transcript isoform diversity across tissues, we analyzed the Cap Analysis of Gene Expression (CAGE) data from the *FANTOM* consortium^6^. These data provide genome-wide quantitative information of transcriptional start sites (TSS) for many cell types. For each subset of the *GTEx* data, we generated a subset of *FANTOM* samples with the same composition of cell types (as long as the samples existed and had replicates). For instance, based on the cell types from *subset A* of the *GTEx* data, we selected a set of *FANTOM* samples consisting of caudates, cerebellums, cortexes, hippocampus and putamens. Then, for each of the three subsets of the *FANTOM* data, we tested each gene for changes in the relative usage of alternative TSS across cell types. At a false discovery rate of 10%, we found 2,402, 6,763 and 2,778 genes with tissue-dependent usage of TSS across subsets *A*, *B* and *C*, respectively. Furthermore, the three lists of genes with differential TSS usage were in very good agreement with the counterpart lists of genes with TDU from the *GTEx* subsets (Table S6). When considering the genes with differential TSS usage across cell types, 79% (1,904) of *subset A*, 80% (5,427) of *subset B* and 60% (1,657) of *subset C* also showed transcript isoform regulation in the corresponding *GTEx* subsets.

Figure 4 shows three examples of genes with TDU patterns that were explained by the usage of alternative TSS. In *subset A*, we found that the gene *Growth Arrest Specific 7* (*GAS7*) expressed tissue-specific isoforms (Figure S8). From the coverage of sequenced RNA fragments along the genome, we suspected that transcription initiated more upstream in cerebellum compared to cerebral cortex. The CAGE data revealed 5 major clusters of TSS for *GAS7*, of which 2 were strongly used in cerebellum and were practically absent from cerebral cortex (Figure 4A). The differential usage of these 2 TSS clusters explained the upstream transcription seen in cerebellum that was not observed in cerebral cortex. Similarly, by exploring the data for the gene *Keratin 8* (*KRT8*) in *subset B*, we found patterns of TDU that were very prominent in thyroid tissue compared to subcutaneous adipose tissue (Figure S9). These patterns of TDU were explained by the usage of a TSS located in the middle of the gene body that resulted in the expression of shorter transcript isoforms. This internal TSS of *KRT8* was used very frequently in thyroid tissue and was absent in subcutaneous adipose tissue (Figure 4B). We found the exact same pattern for the gene *Nebulette* (*NEBL*) in *subset C* of the data. For this gene, the usage of an internal TSS resulted in transcript isoforms that excluded several 5’ exons. This internal TSS was used very frequently in heart tissue, while it was absent in pancreas tissue (Figure 4C, Figure S10).

**Figure 4:**
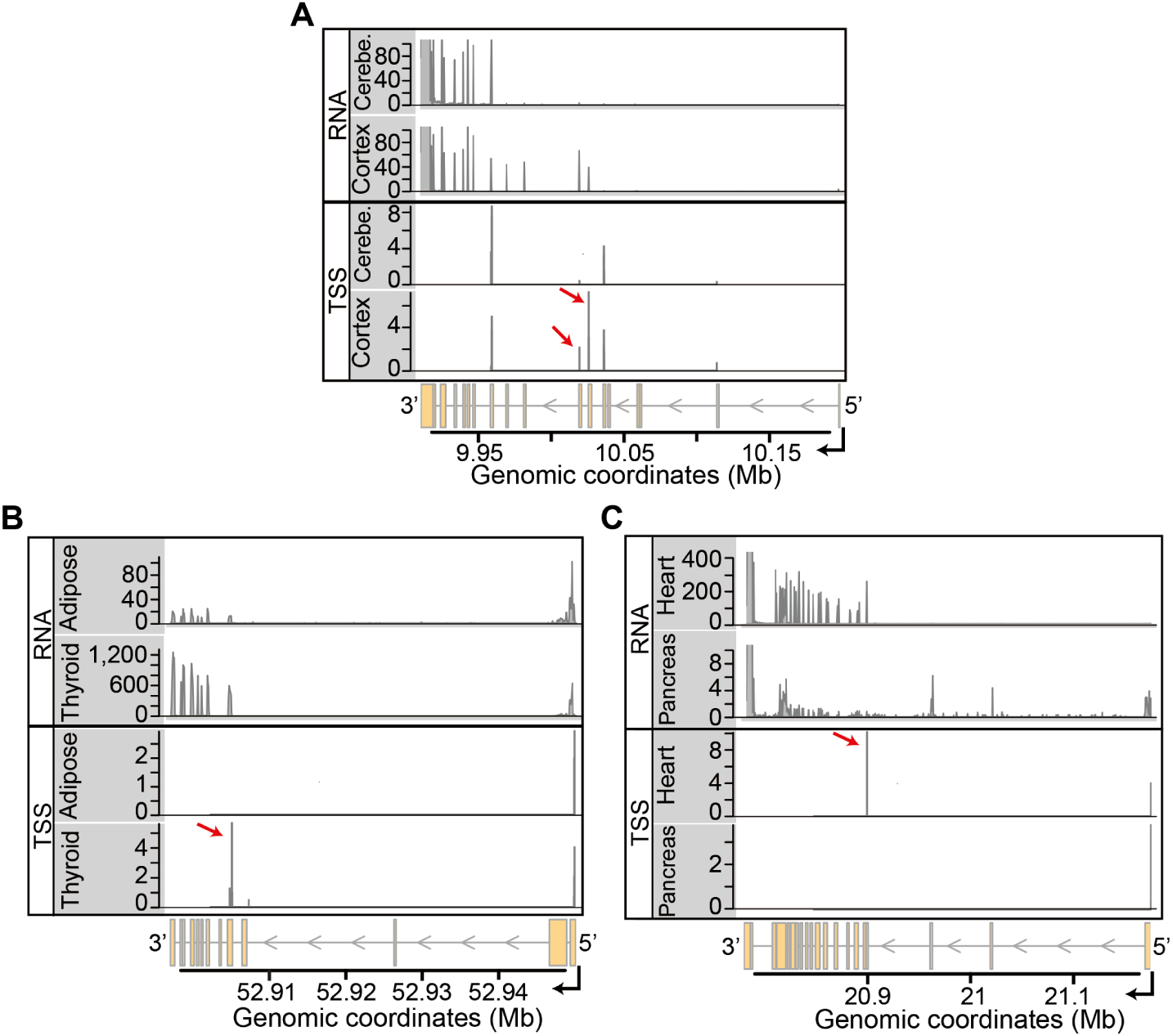
Integration of RNA-seq and CAGE data. Each panel displays an example of a gene where the usage of alternative transcription start sites explains the patterns of TDU. **(A)** Coverage tracks (*y*-axes) of RNA-seq and CAGE data for cerebral cortex and cerebellum are shown along the genomic coordinates (*x*-axis) of the locus of gene *GAS7*, located on chromosome 17. The upper two tracks show RNA-seq data from individual *12ZZX*. The lower two tracks show mean CAGE counts (on *log_2_* scale) for each annotated TSS. Cortex uses two transcription start site clusters (pointed by the red arrows) that are absent in cerebellum. The differential usage of these two TSS explains the upstream RNA-seq coverage seen in cortex. **(B)** Same as in Figure 4A, but showing data of thyroid and subcutaneous adipose tissue along the genomic coordinates of the *KRT8* locus on chromosome 12. The RNA-seq data corresponds to the individual *11EI6*. The internal TSS cluster that is indicated by the red arrow is strongly used in thyroid tissue, resulting in the expression of short transcript isoforms. **(C)** Same as in Figure 4A, but showing data of heart and pancreas along the genomic coordinates of the *NEBL* locus on chromosome 10. The RNA-seq data corresponds to the individual *ZF29*. In heart, the usage of an internal TSS (indicated by the red arrow) results in the expression of transcript isoforms that exclude several 5’ exons of the gene.

Our integrative analysis of two orthogonal sources of data (independent samples, different technologies) confirms that there is an abundance of alternative TSSs that are used in a tissue-dependent manner and that result in TDU.

### Tissue-dependent splicing of protein-coding exons is rare

We asked which regions of genes were subject to tissue-dependent exon usage. We integrated information from the ENSEMBL and APPRIS databases to annotate each exonic region. Importantly, APPRIS uses information about protein structures, functional data and cross-species conservation to infer which transcript isoforms are likely to code for functional proteins. APPRIS flags transcript isoforms that are predicted to code for functional proteins as principal isoforms, while the rest of the transcripts are marked as non-principal isoforms^17^. Using these sources of information, we classified each exonic region into 5 categories: (1) exonic regions coding for principal isoforms, (2) exonic regions coding only for non-principal isoforms, (3) 5’ untranslated exonic regions (5’ UTR), (4) 3’ untranslated exonic regions (3’ UTR), and (5) untranslated exons belonging to non-coding processed transcripts. Then, for each subset of the *GTEx* data, we generated a background set of exons with the same distributions of both mean counts and exon widths as the set of exonic regions with TDU.

We found that the proportions among the 5 exon categories were very different between exonic regions with tissue-dependent usage due to alternative splicing (TDUAS), exonic regions with TDU but no evidence of alternative splicing (TDU-NAS) and the background sets of exons (*p*-value < 2.2.10^−16^, *χ*^2^-test, Figure 5A, Figure S11 and Table S7). Specifically, exonic regions with TDU-AS were depleted among those coding for principal isoforms and enriched among exonic regions coding for non-principal isoforms and 3’ UTRs. Our analysis also revealed that exons from non-coding processed transcripts, despite being weakly expressed, were alternatively spliced very frequently in a tissue-dependent manner (Figure 5, Figure S11 and Figure S12).

**Figure 5:**
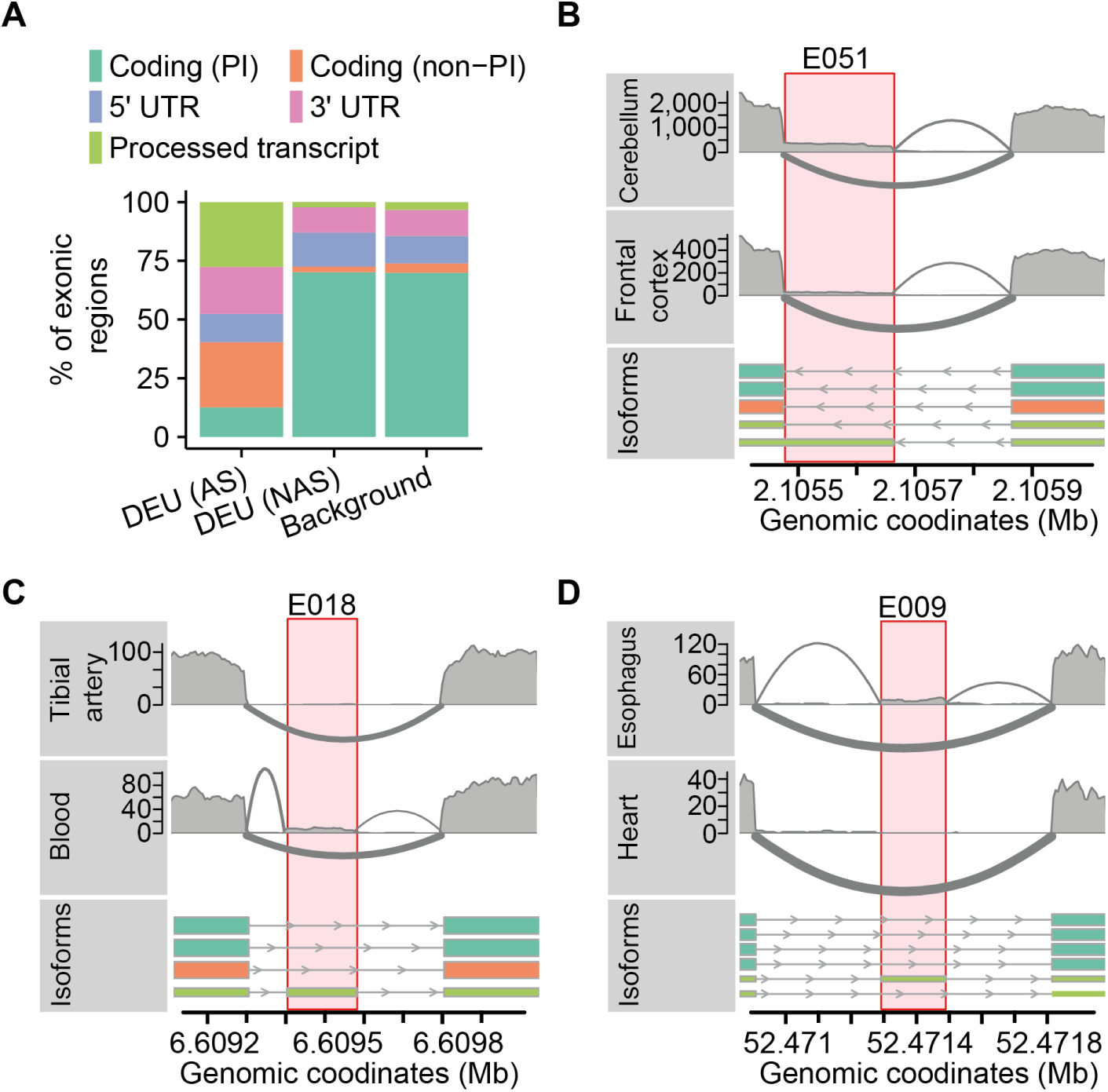
Alternative splicing is infrequent among coding exons. **(A)** The percentage of exonic regions (*y*-axis) is shown for three subsets of exons: (1) exonic regions with TDU due to alternative splicing [DEU (AS)], (2) exonic regions with TDU without evidence of alternative splicing [DEU (NAS)] and (3) a background set of exons matched for expression and exon width. Each color represents a different category of exons according to transcript biotypes: exons coding for principal transcript isoforms [Coding (PI)], exons coding for non-principal transcript isoforms [Coding (non-PI)], 5’ UTRs, 3’ UTRs and exons from non-coding processed transcripts [Processed transcripts]. **(B)** Sashimi plot representation of the RNA-seq data from frontal cortex and cerebellum of individual *WL46*. The lower data track shows the transcript isoforms of the gene *PKD1*. The transcripts are colored according to their biotype (the color legend is the same as in Figure 5A). The highlighted exon (*E051*) belongs to a non-coding transcript and is differentially spliced across tissues. **(C)** Same as in Figure 5B, but showing data from tibial artery and whole blood of the individual *ZTPG*. Transcripts from the gene *MAN2B2* along chromosome 4 are shown. The highlighted exon (*E018*) belongs to a non-coding transcript and is differentially spliced across tissues. **(D)** Same as in Figure 5B, but showing data from esophagus tissue (muscularis) and heart tissue (left ventricle) of the individual *111YS*. The lower track shows the transcripts annotated for gene *NISCH* along chromosome 4. The highlighted exon (*E009*) belongs to a non-coding transcript and is differentially spliced across tissues.

We also found that exonic regions with TDU-NAS showed a slight yet significant enrichment among 5’ UTR exons compared to the background (*p*-value < 1.2 · 10^−7^, *χ*^2^-test, Figure 5A and Figure S11). TDU-NAS occurred frequently among 3’UTR regions compared to the background (*p*-value < 1.2 · 10^−7^, *χ*^2^-test). However, this enrichment among 3’UTR regions was observed only in subsets *B* and *C* of the *GTEx* data (Figure S11).

## DISCUSSION

Our analysis highlights two aspects of transcript isoform diversity that have been under-appreciated. First, alternative splicing is not the main process by which transcript isoform diversity is regulated across tissues. Instead, most recurrent transcript isoform differences are consistent with the alternative usage of transcription start and polyadenylation sites. Second, most of the splicing that is regulated differentially across tissues affects untranslated exons and therefore may not have direct consequences on the protein products.

We analyzed 798 human transcriptomes covering 23 different cell types. This large and comprehensive dataset together with the analytical approach illustrated in Figure 1A enabled us to identify alternative transcription initiation and polyadenylation sites as the principal sources of transcript isoform differences across human cell types. It has been suggested that the regulation of gene expression levels is the main driver of cell type specificity, with splicing playing a complementary role^36^. Our analysis implies that alternative transcription initiation and polyadenylation sites make a sizeable contribution to cellular phenotypes in normal human physiology, and that this contribution is, in some measures, larger than that of splicing.

Transcriptome-wide studies have shown that, in a given cell type, most genes express one major isoform at high levels, while the rest of the isoforms (i.e. the minor isoforms) are expressed at lower levels^37,7^. Importantly, protein isoforms detected in large-scale proteomic experiments are consistent with both the major RNA isoforms and the principal isoforms from the APPRIS database^18^. By combining these lines of evidence with our results, it is strongly suggestive that most tissue-dependent splicing does not have consequences at the proteome level. We support this thesis with three main sources of evidence:

- First, tissue-dependent splicing is enriched among untranslated exons, particularly among exons from non-coding transcript isoforms.
- Second, tissue-specific splicing is depleted among exons coding for principal protein isoforms.
- Third, the exon categories where tissue-dependent splicing is more common are weakly expressed. Thus, the patterns of tissue-specific splicing are explained by tissue-specific expression of minor transcript isoforms.

The remaining open question is, if tissue-dependent splicing has little effect at the proteome level, what are its functions, if any, at the transcriptome level? A parsimonious answer may involve post-transcriptional regulation. Recent CRISPR-mediated interference screens identified 499 long non-coding RNAs that were essential for cell growth, of which 89% of these showed growth-modifying phenotypes that were exclusive to one cell type^38^. Similar screens at the transcript isoform level would be helpful to evaluate the essentiality of the thousands of non-coding and non-translated tissue-specific isoforms derived from protein-coding loci.

Alternative usage of promoters, alternative splicing of exons and alternative usage of polyadenylation sites are highly interleaved^1,39^. In the light of our results, it is conceivable that such coordination of decisions on the usage of transcription start sites, alternative exons and transcriptional termination is often cell type specific. The understanding of such coordination will be facilitated by the development of sequencing technologies that provide quantitative measurements of full-length transcripts^40,41,42^. Furthermore, while alternative splicing may have limited effects for protein products, it remains to be seen to what extent tissue-dependent choice of alternative start and termination sites results in truncated versions of proteins.

## Methods

### Data processing and sample selection

We downloaded and decrypted the *GTEx* samples using the *Short Read Archive Toolkit* software. We used genomic and annotation files of the human reference genome version *GRCh38* as provided by the release number 68 of *ENSEMBL*^28^. To avoid any mapping bias, we standardized the read length of all samples. Since most samples consisted of sequenced fragments of 76 bp, we trimmed the sequenced fragments to 76 bp for samples with longer read lengths and excluded the samples with read lengths below 76 bp. Next, we mapped the resulting sequencing fragments to the human reference genome using *STAR v2.4.2a*^29^. We provided the aligner with annotated exon-exon junctions and followed the recommended “2-pass alignment” pipeline to optimize mapping accuracy. We excluded the samples with less than 1,000,000 sequenced fragments as well as those samples where less that 60% of the sequenced fragments could be assigned to a unique position in the reference genome. Since the *GTEx* data did not contain the samples for all tissues of each individual, we defined three large subsets of samples that would enable us to analyze each subset as a fully-crossed design (containing all tissue-individual combinations) while at the same time keeping as many different individuals and tissues as possible. A description of these subsets, which comprised a total of 798 samples, can be found in the main text.

Based on the transcript isoform annotations, we defined reduced models with non-overlapping exonic regions using the *HTSeq*^43^ python scripts from the *DEXSeq* package. Importantly, reduced gene models enabled us to unambiguously assign sequenced fragments to exonic regions. For each of the 798 samples, we tabulated the sequenced fragments to each exonic region. Only reads mapping uniquely to the reference genome were considered for further analysis.

### Relative exon usage coefficients

We model the counts using generalized models of the Negative Binomial family for each subset of the *GTEx* data^30,31^. We denote *k*_*ij*1_ as the number of sequenced fragments mapping to exonic region *i* in sample *j*. When estimating *REUCs*, *k*_*ij*0_ denotes the sum of sequenced fragments mapping to exonic regions of the same gene as exonic region *i* but excluding exonic region *i* (Figure 1B). *k_ij_*_0_ and *k_ij_*_1_ are realizations of a random variable *K_ijl_* that is assumed to follow a Negative Binomial distribution, 
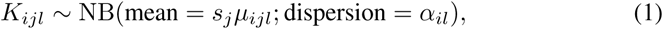

where *s_j_* is a scaling factor that accounts for between-sample differences in sequencing depth and *α_il_* is the dispersion parameter that describes the spread of the count data distribution. *s_ij_* is estimated using the *DESeq* method^44^ and *α_il_* is estimated as in the *DEXSeq* method^30^. The mean *μ_ijl_* is predicted by the model
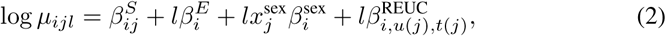

where *l* = 1 when referring to the exonic region *i* and *l* = 0 when referring to the counts from the rest of the exons of the same gene. The coefficients of the model are explained as follows:

- 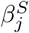 estimates overall gene expression effects on sample *j*.
- Since 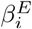 is only included when *l* = 1, it estimates the mean across samples of the logarithmic ratio between the counts from exon *i* with respect to the counts of the rest of the exons of the same gene (i.e. *K*_*ij*1_/*K*_*ij*0_). Therefore, this coefficient is a measure of the average exon usage across all samples.
- 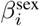 captures sex-dependent differences in exon usage. In the GLM model matrix, *x_ij_* takes the value of −1/2 if sample *j* is from a male individual and 1/2 if sample *j* is from a female individual. Thus, this coefficient estimates the logarithmic fold change of the usage of exonic region *i* for each sex with respect to the average exon usage.
- The *Relative Exon Usage Coefficient* (*REUC*), 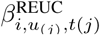, is the interaction coefficient between individual *u*_(*j*)_ and tissue *t*_(*j*)_ from which sample *j* was taken. For exonic region *i*, the coefficient 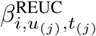 thus estimates the logarithmic fold change in exon usage for each individual-tissue combination with respect to the average exon usage.

The *REUCs* are subjected to a Bayesian shrinkage procedure in order to reduce the mean-variance dependencies ^31,45^.

### Relative spliced-in coefficients

To estimate *Relative Spliced-In Coefficients* (*RSICs*) we use Equation 1 and Equation 2 to model a modified read counting scheme (Figure 1C). *k*_*ij*1_ remains the same as for the *REUCs* fit but *k*_*ij*0_ (i.e. *l* = 0) now denotes the number of sequenced fragments supporting the splicing out from transcripts of exonic region *i* (Figure 1C). For exonic region *i*, the coefficient 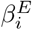 from Equation 2 now measures the mean across samples of the logarithmic ratio between the number of reads supporting the splice in of exonic region *i* and the number of reads supporting the splice out of exonic region *i* (i.e. the average spliced-in (*SI*) coefficient). The coefficient 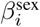 for exonic region i now measures the change of *SI* between each sex with respect to the average *SI*. The *RSIC* for exon *i*, 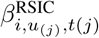 measures the logarithmic fold change in the exon’s *SI* for each tissue-individual combination with respect to the average *SI*. As for the *REUCs*, the *RSICs* are also subjected to the Bayesian shrinkage procedure to eliminate the mean-variance trend ^31^.

Changes in exon usage driven by alternative splicing are reflected in both *REUCs* and *RSICs*. Changes in exon usage due to alternative initiation or termination sites of transcription, which do not result in exon-exon junction reads, are only reflected by *RSICs*.

### Estimation of tissue-dependance score

For each exonic region on each subset of the data, we estimated a score based on the *REUCs* to measure to what extent the usage of each exonic region was tissue-dependent. First, the *REUCs* of a given exonic region *i* were expressed as the number of standard deviations away from the median of the exon’s *REUCs*,
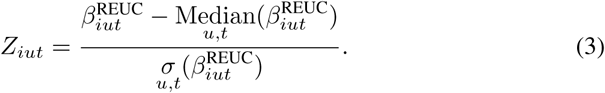

Then, the tissue-dependence score for exonic region *i* was defined by
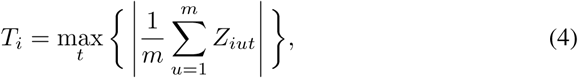

with *m* being the number of individuals on the data subset.

### Analysis of variance of *REUCs* and *RSICs*

For each exonic region on each subset of the data, we fitted an analysis of variance model,
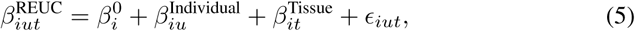

using ordinary least squares regression to minimize the residual sum of squares (RSS),
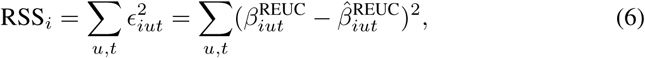

where 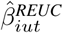 are the *REUC* values predicted by the model. In order to estimate the coefficient of partial determination (*R*^2^) for the tissue predictor (i.e. the proportion of total variance that can be attributed to variance across tissues), we fitted a reduced model lacking the 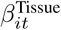 term,
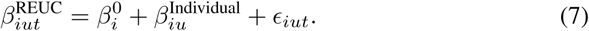

The *R*^2^ for a given exon i is then calculated by,
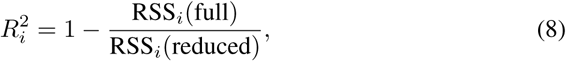

where RSS*_i_* (full) is the RSS from the full model (i.e. Equation 5) and RSS*_i_* (reduced) is the RSS from the reduced model (i.e. Equation 7). The same procedure was followed to estimate *R*^2^ on the *RSICs* but using 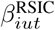 as the response variable in Equation 5 and in Equation 7.

### Genomic analysis

To test for over-representation of features among the genes with TDU, we used the *R CRAN* package *MatchIt*^46^ to generate background sets of genes with the same distribution of expression strength and number of exonic regions as the genes with TDU. Genes were classified according to *ENSEMBL* annotations and we used a *χ*^2^-test for differences between genes with TDU and the background set of genes. Gene biotypes were retrieved from *ENSEMBL* using the *Bioconductor*^47^ package *biomaRt*^48^. For enrichment of features among exons with tissue-dependent usage (*TDU*), we also used *MatchIt* to generate background sets of exons with the same distribution of expression strength and exon widths. We tested for differences between exons with *TDU* and the background set of exons using a *χ*^2^-test.

Operations on genomic ranges were done using the *Bioconductor* package *GenomicRanges*^49^. Data visualizations and graphics were generated using the *Bioconductor* packages *ggplot2*^50^ and *Gviz*^51^.

### Data accesibility and reproducibility

The raw data used for this project is part of the *GTEx* Project and can be found in dbGAP under the accession identifier phs000424.v6.p1. The package *HumanTissues-DEU* contains the *R* data objects and code needed to reproduce the analysis and figures presented in this manuscript.

## Acknowledgements

We would like to thank Alvis Brazma and Vicent Pelechano for critical reading of this manuscript. We thank the *GTEx* and *FANTOM* consortiums for providing access to their data. This work is part of the SOUND Project funded by the H2020 Programme of the European Commission (grant agreement 633974). AR would like to acknowledge funding from the National Institutes of Health (grant R01HG005220).

